# A rapid and simple method for assessing and representing genome sequence relatedness

**DOI:** 10.1101/569640

**Authors:** M Briand, M Bouzid, G Hunault, M Legeay, M Fischer-Le Saux, M Barret

## Abstract

Coherent genomic groups are frequently used as a proxy for bacterial species delineation through computation of overall genome relatedness indices (OGRI). Average nucleotide identity (ANI) is a widely employed method for estimating relatedness between genomic sequences. However, pairwise comparisons of genome sequences based on ANI is relatively computationally intensive and therefore precludes analyses of large datasets composed of thousands of genome sequences.

In this work we proposed a workflow to compute and visualize relationships between genomic sequences. A dataset containing more than 3,500 Pseudomonas genome sequences was successfully classified with an alternative OGRI based on k-mer counts in few hours with the same precision as ANI. A new visualization method based on zoomable circle packing was employed for assessing relationships among the 350 groups generated. Amendment of databases with these Pseudomonas groups greatly improved the classification of metagenomic read sets with k-mer-based classifier.

The developed workflow was integrated in the user-friendly KI-S tool that is available at the following address: https://iris.angers.inra.fr/galaxypub-cfbp.

## Introduction

Species is a unit of biological diversity. Prokaryotic species delineation historically relies on a polyphasic approach based on a range of genotypic, phenotypic and chemo-taxonomic (e.g. fatty acid profiles) data of cultured specimens. According to the List of Prokaryotic Names with Standing in Nomenclature (LPSN), approximately 15,500 bacterial species names have been currently validated within this theoretical framework [1]. According to different estimates the number of bacterial species inhabiting planet Earth is predicted to range between 107 to 1012 species [2,3], the genomics revolution has the potential to accelerate the pace of species description.

Prokaryotic species are primarily described as cohesive genomic groups and approaches based on similarity of whole genome sequence, also known as overall genome relatedness indices (OGRI), have been proposed for delineating species. Genome Blast Distance Phylogeny (GBDP [4]) and Average nucleotide identity (ANI) are currently the most frequently used OGRI for assessing the relatedness between genomic sequences. Distinct ANI algorithms such as ANI based on BLAST (ANIb [5]), ANI based on MUMmer (ANIm [6]) or ANI based on orthologous genes (OrthoANIb [7]; OrthoANIu [8]; gANI,AF [9]), which differ in their precision but more importantly in their calculation times [8], have been developed. Indeed, improvement of calculation time for whole genomic comparison of large datasets is an essential parameter. As of November 2018, the total number of prokaryotic genome sequences publicly available in the NCBI database is 170,728. Considering an average time of 1 second for calculating ANI values for one pair of genome sequences, it would take approximately 1,000 years to obtain ANI values for all pairwise comparisons.

The number of words of length k (k-mers) shared between read sets [10] or genomic sequences [11] is an alignment-free alternative for assessing the similarities between entities. Methods based on k-mer counts, such as SIMKA [10], Kmer-db [12] or Mash [13] can quickly compute pairwise comparison of multiple metagenome read sets with high accuracy. In addition, specific k-mer profiles are now routinely employed by multiple read classifiers for estimating the taxonomic structure of metagenome read sets [14–16]. While these k-mer based classifiers differ in terms of sensitivity and specificity [17], they rely on accurate genome databases for affiliating a read to a taxonomic rank.

The objective of the current work was to propose a workflow to quickly compute OGRI and visualize the outputs in an efficient manner. An alternative method based on k-mer counts was first evaluated to study species delimitation on extensive genome datasets. We employed k-mer counting to assess the similarity among genome sequences belonging to the Pseudomonas genus. Indeed, this genus contains an important diversity of species (n = 207), whose taxonomic affiliation is under constant evolution [18–24], and numerous genome sequences are available in public databases. We also proposed an original visualization tool based on D3 Zoomable Circle Packing (https://gist.github.com/mbostock/7607535) for assessing the relatedness of thousands of genome sequences. Finally, the benefit of taxonomic curation of reference database on the taxonomic affiliation of metagenomics read sets was assessed. The developed workflow was integrated in the user-friendly KI-S tool which is available in the galaxy toolbox of CIRM-CFBP (https://iris.angers.inra.fr/galaxypub-cfbp).

## Methods

### Genomic dataset

All genome sequences (n=3,623 as of April 2017) from the Pseudomonas genus were downloaded from the NCBI database (https://www.ncbi.nlm.nih.gov/genome/browse#!/overview/).

### Calculation of Overall Genome Relatedness Indices

The percentage of shared k-mers between genome sequences was calculated with Simka version 1.4 [10] with the following parameters (abundance-min 1 and k-mer lengths of 11, 13, 15, 17 and 19). Distances between genomic sequences were also calculated using k-mers based methods including FastANI [25], Kmer-db [12] (with the following parameters -k 15, distance min) and Mash [13]. The percentage of shared k-mer was compared to ANIb values calculated with PYANI version 0.2.3 [26]. Due to the computing time required for ANIb calculation, only a subset of Pseudomonas genomic sequences (n=934) were selected for this comparison. This subset was composed of genome sequences containing less than 150 scaffolds.

### Development of KI-S workflow

An integrative workflow named KI-S (Kinship relationships Identification with Shared k-mers) was developed. At the first step, the percentage of shared k-mers (k can be selected by the user) between genome sequences is calculated with Simka [10] or estimated with Kmer-db [12]. A custom R script is then employed to cluster the genome sequences according to their connected components at different selected thresholds (e.g. 50% of shared 15-mers). The clustering result is visualized with Zoomable Circle Packing representation with the D3.js JavaScript library (https://gist.github.com/mbostock/7607535). The source code of the KI-S workflow is available at the following address: https://sourcesup.renater.fr/wiki/ki-s. A wrapper for accessing KI-S in a user-friendly Galaxy tool is also available at the following address: https://iris.angers.inra.fr/galaxypub-cfbp.

### Taxonomic inference of metagenomic read sets

Classification of nine metagenomic read sets derived from seed, germinating seeds and seedlings of common bean (Phaseolus vulgaris var. Flavert) were estimated with Clark version 1.2.4 [16] at the species level. Clark is a method based on a supervised sequence classification using discriminative k-mers [16]. These metagenomic datasets were selected because of the high relative abundance of reads affiliated to Pseudomonas [27]. The following Clark parameters -k 31 -t <minFreqTarget> 0 and -o <minFreqtObject> 0 were used for the taxonomic profiling. Indeed, reducing k increase the number of read assignments but also increase the probability of misclassification [16]. Three distinct Clark databases were employed: (i) the original Clark database from NCBI/RefSeq (ii) the original Clark database supplemented with the 3,623 Pseudomonas genome sequences and their original NCBI taxonomic affiliation (iii) the original Clark database supplemented with the 3,623 Pseudomonas genome sequences whose taxonomic affiliation was corrected according to the reclassification based on the number of shared k-mers. For this third database, genome sequences were clustered at >50% of 15-mers, which corresponded to the species level.

## Results

### Selection of optimal k-mer size, k-mers indexing softwares and percentage of shared k-mers

Using the percentage of shared k-mers as an OGRI for species delineation first required the determination of the optimal k-mer size. This was performed by comparing the percentage of shared k-mers with SIMKA [10] to a widely employed OGRI, ANIb [5], among 934 Pseudomonas genome sequences. Since the species delineation threshold was initially proposed following the observation of a gap in the distribution of pairwise comparison values [28], the distribution profiles obtained with k-mer lengths 11, 13, 15, 17 and 19 were compared to ANIb values. Short k-mers (k =11) were evenly shared by most strains and therefore non-discriminatory (**Fig. 1**). As the length of the k-mer increased (k ≥ 13), a multimodal distribution based on several peaks was observed (**Fig. 1**). This multimodal distribution reflects a genetic discontinuity previously observed in several studies [6, 9, 25]. For example, for k=15, the first peak observed (at 15% of shared 15-mers) is related to the genome sequences that do not belong to the same species. Then, the peaks at 50 and 80% shared 15-mers corresponded to genome sequences associated to the same species and subspecies, respectively. The last peak at 100% of shared k-mers was related to identical genome sequences. Since increasing k-mer lengths beyond 15 did not improve the resolution of the multimodal distribution but leads to a more rapid drop in the percentage of shared k-mers between strains, we selected k=15 for subsequent analyses.

**Figure 1:**
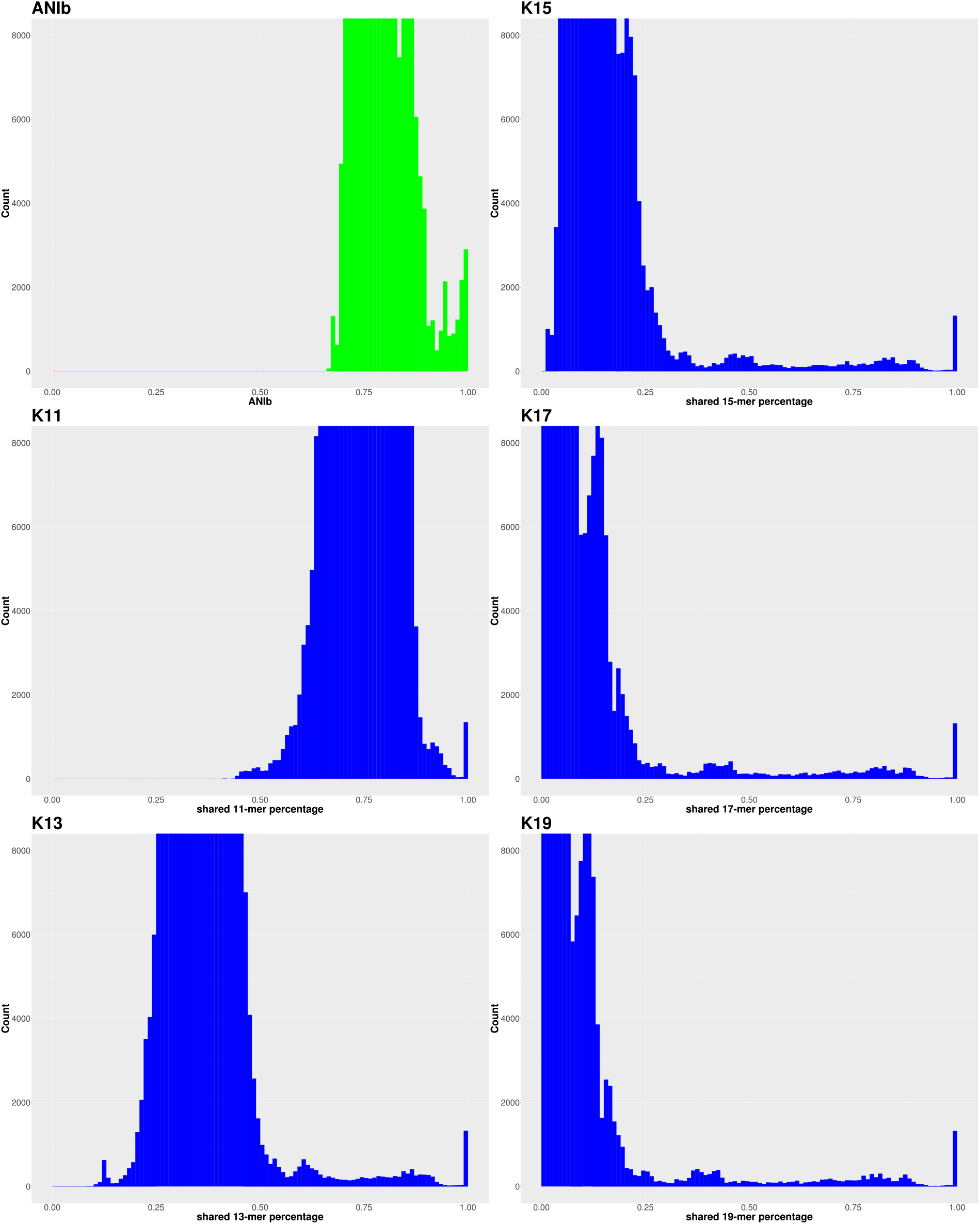
Distribution of shared k-mers values. Relatedness between genome sequences were estimated with ANIb (green) or shared k-mers (blue). The x axis represents ANIb or percentage of shared k-mers while the y axis represents the number of values by class in the subset of 934 Pseudomonas genomic comparison.

Several tools using k-mers for estimating genomes relatedness (e.g. FastANI, Simka, Kmer-db, Mash) were available at the time of analysis. Mash outperformed all others tools in term of computation time (**Table 1**).

**Table 1:** Comparison of softwares that estimate genome relatedness.

| Software | Time | RAM | Reference |
| --- | --- | --- | --- |
| PYANI | 3 months | ND | 26 |
| FastANI | 11 days | 15 Go | 25 |
| Simka | 4 hours | 18 Go | 10 |
| Kmer-db | 40 minutes | 25 Go | 12 |
| Mash | 7 minutes | 26 Mo | 13 |

Indeed, computation time of 15-mers for 934 genome sequences was 7 min with Mash on a DELL Power Edge R510 server, while it took approximately 3 months for obtaining all ANIb pairwise comparisons. The outputs of these different softwares were next compared to ANIb (**Fig. 2**). FastANI was the best estimator of OGRI as indicated by the strong linear relationship of average pairwise similarities with ANIb values (**Fig. 2**). However, one small caveat is that sequence similarity values were ignored by FastANI when below 0.76 value, which artificially improved the linear relationship. According to these linear relationships, Simka and Kmer-db performed reasonably well for ANIb values above 0.9, while MASH was restricted to ANIb values above 0.95. Hence, Kmer-db and Simka were selected in KI-S workflow since these tools are the best compromise between quick computation time and robust genome relatedness indexes (**Table 1** and **Fig. 2**).

**Figure 2:**
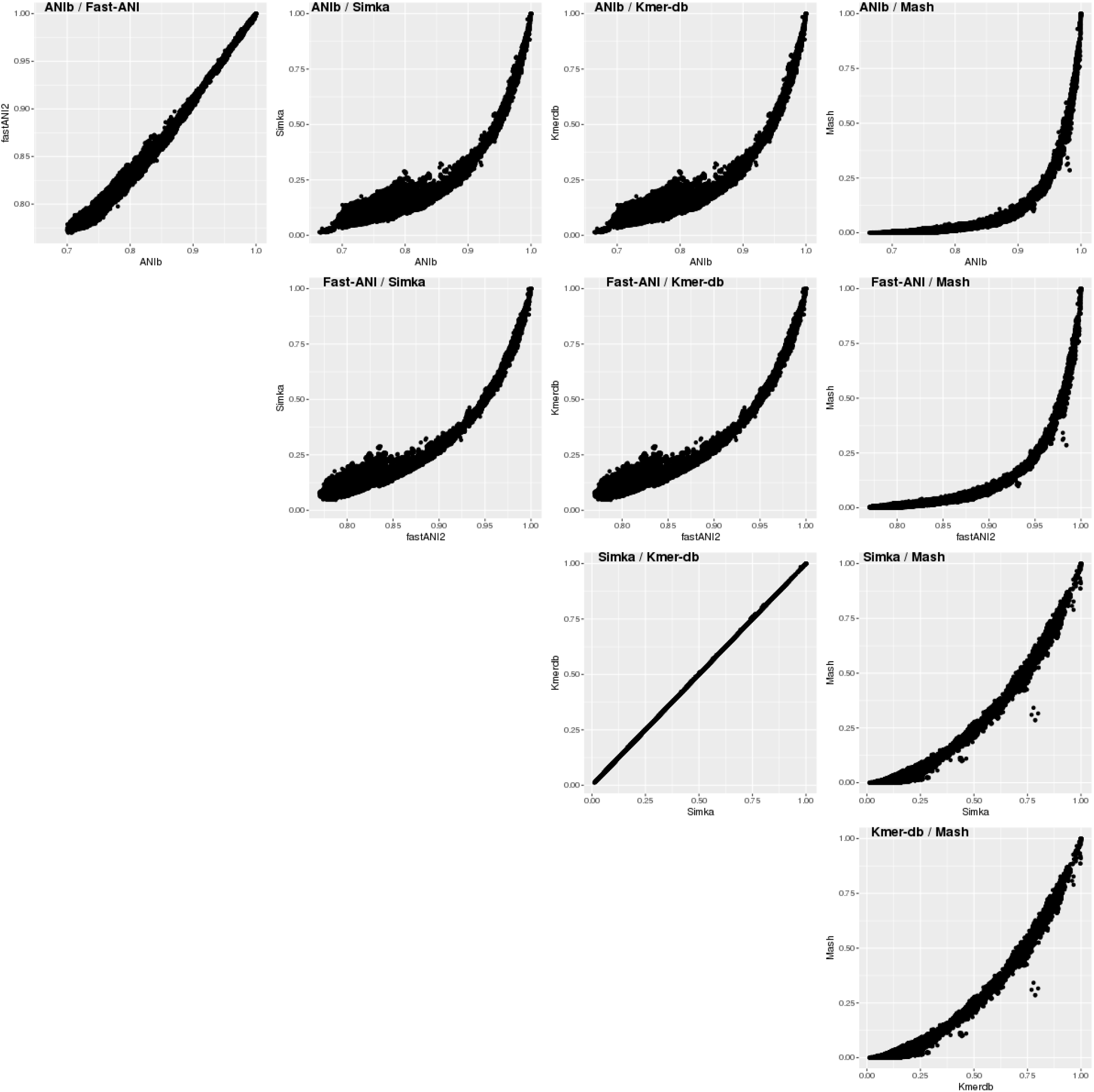
Comparison of average pairwise similarity between genomes sequences. Overall genome relatedness indexes of 934 Pseudomonas genomes sequences were calculated with PYANI [26] and four different k-mers indexing softwares: FastANI [25], Simka [10], Kmer-db [12] and MASH [13].

We next investigated what percentage of shared 15-mers corresponds to an ANIb value of 0.95, a threshold commonly employed for delineating bacterial species [5]. Fifty percent of 15-mers were closed to an ANIb value of 0.95 (**Fig. 2**). More precisely, the median percentage of shared 15-mers was 49% [34%-66%] for ANIb value ranging from 0.94 to 0.96. The 934 genomic sequences were clustered in 329 and 315 groups at an ANIb value of 0.95 and 50% of 15-mers, respectively. The composition of these groups was identical between the two approaches for 302 groups that contained 808 genomic sequences. The 27 additional groups obtained with ANIb were nestled within the 13 additional groups derived from 50% of shared 15-mers.

### Classification of Pseudomonas genome sequences

The percentage of shared 15-mers was then used to investigate the relatedness between 3,623 Pseudomonas publicly available genome sequences. At a threshold of 50% of 15-mers, we identified 350 groups. The group containing the most abundant number of genome sequences was related to P. aeruginosa (n = 2,341), followed by the phylogroups PG1 (n = 111), PG3 (n = 92) and PG2 (n = 74) of P. syringae species complex ([19]; **Table S1**). At the clustering threshold employed, 185 groups were composed of a single genome sequence, therefore highlighting the high Pseudomonas strain diversity. Graphical representation of-clustering by dendrogram for a large dataset is generally not optimal. Here we employed Zoomable circle packing as an alternative to dendrogram for representing similarity between genome sequences (**Fig. 3** and **FigS1.html**). The different clustering thresholds that can be superimposed on the same graphical representation allow the investigation of inter- and intra-groups relationships (**Fig. 3** and **FigS1.html**).

**Figure 3:**
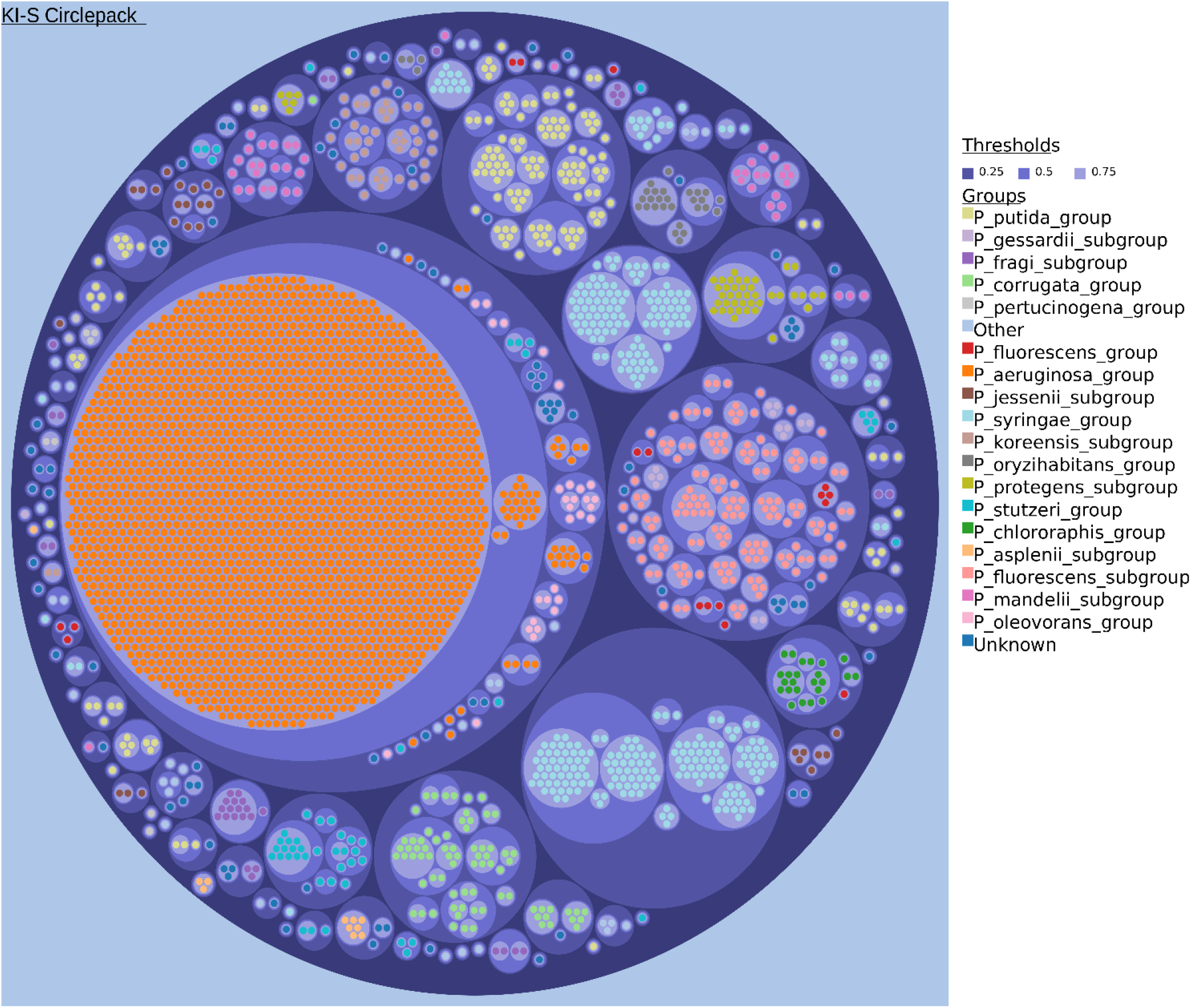
Clustering of Pseudomonas genome sequences. Zoomable circle packing representation of Pseudomonas genome sequences (n = 3,623). Similarities between genome sequences were assessed by comparing the percentage of shared 15-mers. Each dot represents a genome sequence, which is colored according to its group of species [19,24]. These genome sequences have been grouped at three distinct thresholds for assessing infraspecific (0.75), species-specific (0.5) and interspecies relationships (0.25).

### Improvement of taxonomic affiliation of metagenomic read sets

The taxonomic composition of metagenome read sets is frequently estimated with k-mer based classifiers. While these k-mer based classifiers differ in terms of sensitivity and specificity, they all rely on accurate genome databases for affiliating a read to taxonomic rank. Here, we investigated the impact of database content and curation on taxonomic affiliation. Using Clark [16] as a taxonomic profiler with the original Clark database, we classified metagenome read sets derived from bean seeds, germinating seeds and seedlings [27]. Adding the 3,623 Pseudomonas genome sequences with their original taxonomic affiliation from NCBI to the original Clark database did not increase the percentage of classified reads (**Fig. 4**). However, adding the same genome sequences reclassified in groups according to their percentage of shared k-mers (k=15; threshold= 50%) increased 1.4-fold on average the number of classified reads (**Fig. 4**).

**Figure 4:**
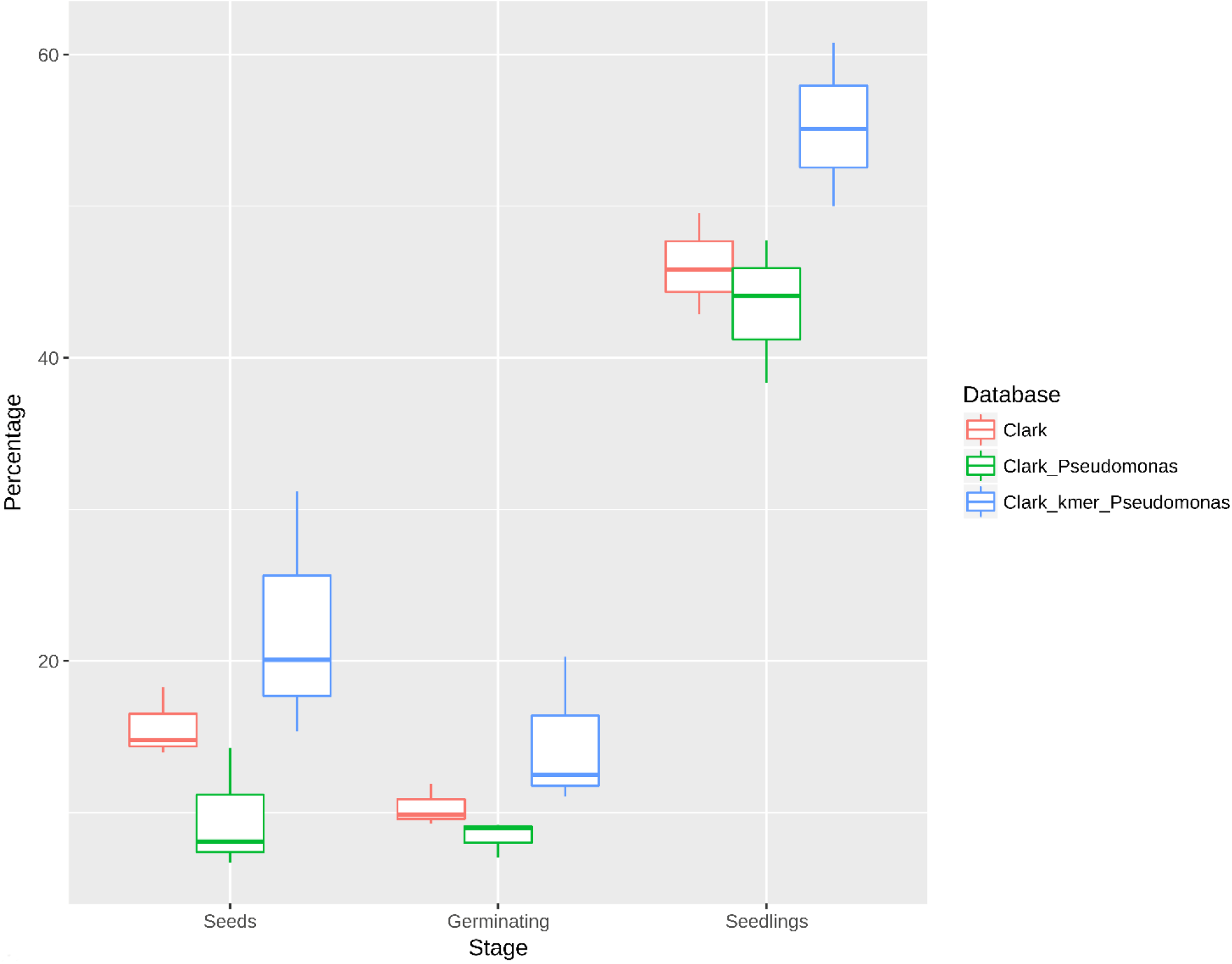
Percentage of classified reads. Classification of metagenome read sets derived from bean seeds, germinating seeds and seedlings with Clark [16]. Three distinct databases were employed for read classification: the original Clark database (red), Clark database supplemented with 3,623 Pseudomonas genome sequences (green) and the Clark database supplemented with 3,623 Pseudomonas genome sequences that were classified according to their percentage of shared k-mers (blue).

## Discussion

Classification of bacterial strains on the basis on their genome sequence similarities has emerged over the last decade as an alternative to the cumbersome DNA-DNA hybridizations [4, 29]. Although ANIb is one widely employed method for investigating genomic relatedness, its intensive computational time prohibits its use for comparing large genome datasets [8]. In contrast, investigating the percentage of shared k-mer is scalable for comparing thousands of genome sequences.

In a method based on k-mer counts, choosing the length of k is a compromise between accuracy and speed. The distribution of shared k-mer values between genome sequences is impacted by k length. For k = 15, four peaks were observed at 15%, 50%, 80% and 100% of shared k-mers. The second peak is close to an ANIb value of 0.95 and falls in the so called grey or fuzzy zone [29] where taxonomists might decide to split or merge species. Hence, according to our working dataset, it seems that 50% of 15-mers is a good proxy for estimating Pseudomonas group. Despite the diverse range of habitats colonized by different Pseudomonas populations [22], it is likely that the percentage of shared k-mers has to be adapted when investigating other bacterial genera. Indeed, since population dynamics, lifestyle and location impact molecular evolution, it is somewhat illusory to define a fixed threshold for species delineation [30]. While 15-mers is a good starting point for investigating infra-specific to infra-generic relationships between genome sequences, the computational speed of KI-S offers the possibility to perform large scale genomic comparisons at different k sizes to select the most appropriate threshold.

Genomic relatedness using whole genome sequences has become the standard method for bacterial strain identification and bacterial taxonomy [4,29,31]. This is primarily motivated by fast and inexpensive sequencing of bacterial genomes together with the limited availability of cultured specimen for performing classical polyphasic approach. Whether full genome sequences should represent the basis of taxonomic classification is an ongoing debate between systematicians [32]. While this consideration is well beyond the objectives of this work, obtaining a classification of bacterial genome sequences into coherent groups is of general interest. Indeed, the number of misidentified genome sequences is exponentially growing in public databases. A number of initiatives such as Digital Protologue Database (DPD [33]), Microbial Genomes Atlas (MiGA [34]), Life Identification Numbers database (LINbase [35]) or the Genome Taxonomy Database (GTDB [31]) proposed services to classify and rename bacterial strains based on ANIb values or single copy marker proteins. Using the percentage of shared k-mers between unknown bacterial genome sequences and reference genome sequences associated to these databases could provide a rapid complementary approach for bacterial classification. Moreover, KI-S includes a friendly visualization interface that could help systematicians to curate whole genome databases. Indeed, zoomable circle packing could be employed for highlighting (i) misidentified strains, (ii) bacterial taxa that possess representative type strains or (iii) bacterial taxa that contain few genome sequences.

Association between a taxonomic group and its distribution across a range of habitats is useful for inferring the role of this taxa on its host or environment. For instance, community profiling approaches based on molecular marker such as hypervariable regions of 16S rRNA gene have been helpful for highlighting correlations between host fitness and microbiome composition. Higher taxonomic resolution of microbiome composition could be achieved with metagenomics through k-mer based classification of reads. Using the Pseudomonas genus as a use-case, we showed that increasing the breadth of genomic database without investigating the relatedness of genome sequences did not improved the proportion of classified reads. Worse, an unresolved classification may limit the number of species-specific k-mers identified by CLARK and therefore the number of classified reads. Interestingly, an inverse relationship between the number of genome sequences in NCBI RefSeq database and the number of classified reads at the species level was also recently highlighted with other k-mer-based read classifiers [36]. On the contrary, prior classification of the genomic database improve the number of classified reads at the species level. Hence, investigating the relationships between bacterial genome sequences not only benefits bacterial taxonomy but also indirectly microbial ecology.

## Data accessibility

Data are available online: https://www.ncbi.nlm.nih.gov/genome/

## Supplementary material

Script and codes are available online: https://sourcesup.renater.fr/wiki/ki-s/

**TableS1.csv: Pseudomonas groups.** Description of the 350 groups obtained after clustering at 50% of shared 15-mers. For each group, the Pseudomonas group [24] and subgroup [19,24] are displayed.

**FigureS1.html: Zoomable circle packing representation of Pseudomonas genome sequences.** Similarities between genome sequences were assessed by comparing the percentage of shared 15-mers. Each dot represents a genome sequence, which is colored according to its group of species [19,24]. These genome sequences have been grouped at three distinct thresholds for assessing infraspecific (0.75), species-specific (0.5) and interspecies relationships (0.25).

## Acknowledgements

The authors wish to thank Claire Lemaitre and Guillaume Rizk for their assistances with the SIMKA software and Jason Shiller for manuscript assessment and for editing the English.

Version 5 of this preprint has been peer-reviewed and recommended by Peer Community In Genomics (https://doi.org/10.24072/pci.genomics.100001).

